# High-resolution *in vivo* kinematic tracking with injectable fluorescent nanoparticles

**DOI:** 10.1101/2024.09.17.613529

**Authors:** Emine Zeynep Ulutas, Amartya Pradhan, Dorothy Koveal, Jeffrey E. Markowitz

## Abstract

Behavioral quantification is a cornerstone of many neuroscience experiments. Recent advances in motion tracking have streamlined the study of behavior in small laboratory animals and enabled precise movement quantification on fast (millisecond) timescales. This includes markerless keypoint trackers, which utilize deep network systems to label positions of interest on the surface of an animal (*e.g.*, paws, snout, tail, *etc.*). These approaches mark a major technological achievement. However, they have a high error rate relative to motion capture in humans and are yet to be benchmarked against ground truth datasets in mice. Moreover, the extent to which they can be used to track joint or skeletal kinematics remains unclear. As the primary output of the motor system is the activation of muscles that, in turn, exert forces on the skeleton rather than the skin, it is important to establish potential limitations of techniques that rely on surface imaging. This can be accomplished by imaging implanted fiducial markers in freely moving mice. Here, we present a novel tracking method called QD-Pi (<u>Q</u>uantum <u>D</u>ot-based <u>P</u>ose estimation *in vivo*), which employs injectable near-infrared fluorescent nanoparticles (quantum dots, QDs) immobilized on microbeads. We demonstrate that the resulting tags are biocompatible and can be imaged non-invasively using commercially available camera systems when injected into fatty tissue beneath the skin or directly into joints. Using this technique, we accurately capture 3D trajectories of up to ten independent internal positions in freely moving mice over multiple weeks. Finally, we leverage this technique to create a large-scale ground truth dataset for benchmarking and training the next generation of markerless keypoint tracker systems.

## Introduction

Animal movement has traditionally been characterized by manually scoring video footage or direct observation^1–4^. Recent technological advances in machine learning have enabled the study of unrestrained, naturalistic movements with unprecedented speed and accuracy, especially in the field of markerless keypoint tracking^5–10^. State-of-the-art methods such as transfer learning with convolutional networks^9^, U-Net-inspired architectures^10^, and 3D convolution networks^6^ have enabled tracking of discrete computationally-defined points on the surface of laboratory animals trained on a small number of hand-labeled video frames. These advances, in turn, have enabled new methods for classifying animal behaviors^11–13^ and quantifying movement^14,15^.

However, due to variations in experimental setups across laboratories, markerless keypoint trackers still require hand-labeling datasets. This can introduce jitter due to variation between annotators and the inherent ambiguity of labeling certain body parts based solely on surface features, such as positions along the back of a mouse. In rats, where markerless keypoint trackers have been systematically compared against skin-attached fiducials, their predictions were estimated to have precision on the order of ±10 mm^16^, comparable to the distance between many key landmarks on the mouse^17^. This contrasts with commercial motion capture systems used with humans, which have a demonstrated precision of approximately ±0.1 mm^18–20^.

Another limitation of markerless keypoint trackers is that they are trained to identify points on the outside of the animal’s body using surface features visible in videography. As joint and skeletal kinematics can be obscured by soft tissue and fur in rodents^21,22^ and humans^23,24^, it remains unclear if movement of the skeleton can be resolved in this way. This is important because the brain directly controls the muscles, which exert complex forces on the skeleton. It remains unclear if tracking motions on the surface of the skin is a viable strategy for resolving the brain’s control of movement and how it breaks down in disease and injury^21,24–27^.

Though technically demanding, it is possible to directly observe the skeletal system in live rats using X-ray videography, which has demonstrated the principle that skin-derived joint angles and kinematics can greatly diverge from those derived directly from the skeleton^21^. While this technique has very high spatial resolution, X-ray videography is challenging to set up in individual labs and can be limited by radiation dosage, allowing only brief imaging sessions^28–30^.

An alternate strategy for locating landmarks inside the body would be to implant optical tags that can be measured non-invasively. To be successful, this approach must satisfy two important criteria. First, the implanted optical tag must be detectable from outside of the animal. Prior studies suggest that near-infrared I (NIR-I, 650-900 nm) is an ideal spectral window for non-invasive imaging due to minimal light absorption and scattering by skin^31–33^ and hemoglobin^34^. Quantum dots (QDs) are one of the few fluorescent materials in this spectral range that are photostable (*i.e.,* minimal photobleaching) and bright (*i.e.,* high quantum yields and extinction coefficients), making them an attractive material relative to NIR-fluorescent proteins and dyes^32,35–37^. Second, the implanted optical tag should be long-lasting. In addition to being photostable, it is therefore important for the tag to be biocompatible and to have a long half-life *in vivo*. QDs are widely used in the life sciences, and thus are readily available in numerous biocompatible formulations^37–40^ and have been used *in vivo*^41,42^.

Here we present a novel tracking method called QD-Pi (<u>Q</u>uantum <u>D</u>ot-based <u>P</u>ose estimation *in vivo*). QD-Pi uses NIR-I-emitting QDs as injectable, bright, long-lived optical probes that can be imaged non-invasively in freely moving animals using standard machine-vision cameras. We demonstrate that QD-Pi can be used to reliably track keypoint markers inside of the mouse’s body while they freely move in an open plexiglass arena. In proof-of-principle experiments, we show that QDs can be injected subdermally in the subcutaneous fatty tissue and act as bright, long-lived ground truth fiducial markers for keypoint tracking. First, using a simple modified microinjection system, we show that commercially available QDs are sufficient for short-term *in vivo* experiments (up to approximately 72 hours post-injection). Second, we show that microbead-immobilized QDs can be used for long-term *in vivo* experiments (up to approximately 2 weeks post-injection). Then, we describe a large dataset for ground-truthing markerless keypoint trackers, wherein internal QDs and the surface of the body were imaged simultaneously. We used this dataset to benchmark the performance of an existing markerless keypoint tracker, SLEAP^10^, relative to ground-truth fiducials embedded in the skin. We also demonstrate that QDs can be injected intra-articularly into joints for directly tracking joint kinematics. Finally, we show in a proof-of-principle that QDs can be conjugated to antibodies for various applications, including targeting ultra-long-lived proteins to further enhance longevity up to 6 weeks post-injection. Unlike UV ink or reflective piercings^16,43,44^, both of which reflect the movement of the skin, our method represents a plausible path for direct measurement of joints, and potentially muscle and bone as well. Additionally, in contrast with X-ray videography, our method can be scaled to accommodate larger fields of view, does not require expensive equipment, and does not expose animals to radiation. Finally, compared with direct tattooing of the joint^45^, our method is more versatile, does not require major surgery, and has higher contrast. Ultimately, QD-Pi addresses a critical need in the study of motor control in rodents and sets the stage for a generalizable method to track skeletal kinematics in freely moving animals.

## Results

### Quantum dots are viable injectable optical tags in mice

We developed a protocol to inject QDs subdermally into mice to track internal points while they freely move in an arena. We selected QDs as a fluorescent marker because of their brightness, photostability, and broad emission spectral profiles within the NIR-I (650–900 nm) range, which is an ideal window for imaging through skin, fur^31–33^, and blood^34^. QDs are also biocompatible, and their surface chemistry can be readily modified with a variety of biologically functional surface coatings. Here, we utilized QDs with a fluorescence emission peak at 800 nm (referred to as QD800 throughout) coated in polyethylene glycol (PEG), previously demonstrated to be biocompatible when introduced into animals^46^.

We first tested whether QDs could form discrete “tags” without dispersing through the skin and underlying tissue (**Fig. 1a-c**). Ideally, the resulting tag would be large enough to resolve on a standard machine vision camera, yet small enough to mark many different points on the body. As an approximation, we assumed as a basic criterion that fluorescence points should be at least 1 mm in diameter to enable high-resolution tracking (roughly 5 pixels assuming an object distance of 12 inches to the camera based on our camera calibrations).

**Figure 1.**
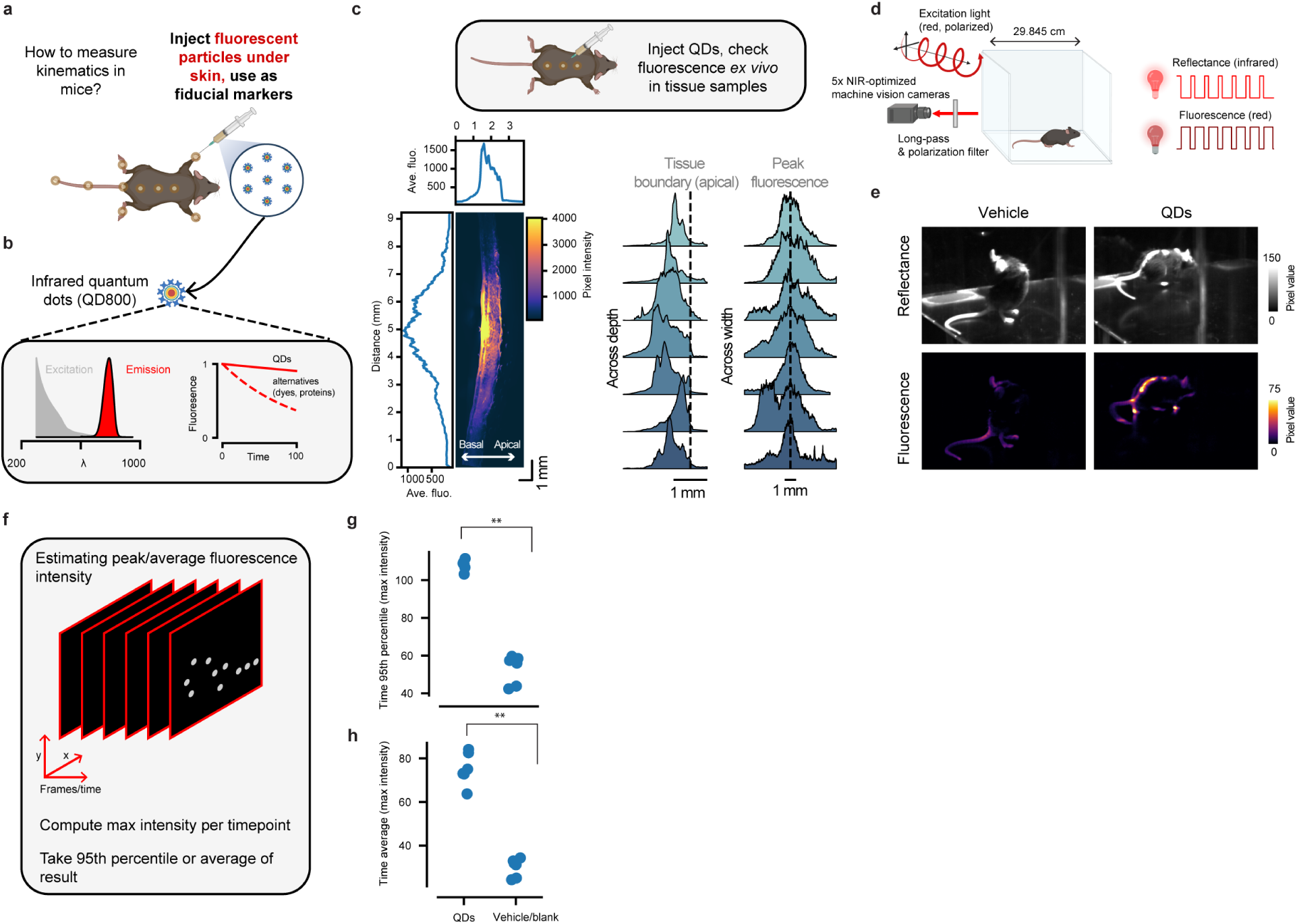
Quantum dots (QDs) can act as optical markers when introduced into the skin. a) Schematic representation of the QD injection procedure. b) Schematic representation of the basic optical properties of NIR-emitting QDs. c) Histological examination of QDs injected into the back of a mouse. Top, schematic of experiment. Injectable QDs were introduced into the skin, after which tissue samples were harvested for imaging. Bottom left, Example fluorescence image of a skin sample after QD injection (QD800.2). Mean projections are shown along the x- and y-axis. Bottom right, fluorescence probability densities at different tissue depths aligned to the tissue boundary (left) or the fluorescence peak (right). Each plot corresponds to a separate injection (QD800.1 or QD800.2). d) Schematic of the plexiglass arena and optical configuration for in vivo imaging. Reflectance and fluorescence images are collected near-simultaneously using IR-emitting LEDs and polarized NIR-I-emitting LEDs, respectively. Five machine vision cameras equipped with long-pass and polarization filters are used to collect images. e) Example reflectance and fluorescence images from vehicle (left) and QD injected mice (QD800.2, right). f) Schematic illustrating how QD fluorescence is measured. First, the max intensity is computed per frame, then either the 95^th^ percentile or the mean is computed across all frames for each mouse and camera. g) 95^th^ percentile pixel intensity comparison of vehicle/blank (n=6 mice) and QD mice (n=6 mice) (p=.002, U=36, f=0, Mann-Whitney U test). Measurements were averaged across the five cameras. h) Average pixel intensity comparison between vehicle/blank or QD injection (p=.002, U=36, f=0, Mann-Whitney U test).

To accomplish this, we developed a simple micro-injection protocol whereby small quantities (<5 µL) of QD800 particles could be injected under the skin of a briefly anesthetized live mouse using a glass micropipette (see **Methods**). We selected QD800 vascular labels, a QD suspension originally designed for visualization of the vasculature (referred to here as QD800.1), for use in initial tests based on their prior use *in vivo* and desirable spectral properties. We then identified three subdermal locations along the back (dorsal midline) for injection and confirmed the size and depth of the fluorescence spots in extracted *ex vivo* tissue sections (**Fig. 1c**). Histology revealed that injection of QD800.1 yielded a bright fluorescent spot in adipose tissue just beneath the dermis, at a depth of 0.72 ± 0.12 mm beneath the skin with a spot diameter of 1.9 ± 0.2 mm (mean ± standard deviation; approximately 10 pixels in images acquired with machine vision cameras equipped with standard 8 mm focal length lenses at a distance of 12 inches, the approximate distance from each camera to the center of the arena in our setup). We then extended our method to 14 internal locations: dorsal and ventral injections to each paw, three injections along the tail, and three injections along the spine (**Fig. 1a**).

In order to test whether QD800.1 could be injected and clearly resolved in freely moving mice, we constructed a plexiglass imaging arena surrounded by five NIR-optimized machine vision cameras, which was an open-top cube with an edge length of 29.845 cm (**Fig. 1d**, **Extended Data Fig. 1a-b**). The cameras were equipped to collect both reflectance images to visualize the location of the mouse and fluorescence images to visualize QD800 fluorescence (730 nm excitation, >800 nm emission). Following optical tag embedding, mice were introduced into the plexiglass arena and were imaged during free behavior over three to five-minute-long sessions (**Fig. 1e, Extended Data Video 1**). Relative to vehicle-injected mice, mice injected with QD800.1 exhibited clearly resolvable fluorescence (signal-to-noise ratio ∼ 2.51 ± 0.25 SD, where we define signal-to-noise ratio as average fluorescence across the whole image per QD-injected mouse relative to average blank/vehicle-injected mice, n=6 QD800 injected mice, composed of n=3 QD800.1 and n=3 QD800.2 mice, and n=6 blank/vehicle-injected mice, **Fig. 1f-h, Extended Data Fig. 2**).

### As free nanoparticles, injected QD signal decays on the timescale of hours in vivo

For our approach to be viable in the context of standard neuroscience experiments (*e.g.,* photometry, imaging, pharmacological or optogenetic manipulations), our optical tags should retain fluorescence for as long as possible — ideally on the order of weeks. To test the longevity of QD800 fluorescence, we monitored fluorescence in freely moving mice across multiple days after subdermal injection (**Fig. 2a-d**). As QD800.1 is a suspension of nanoparticles (10-20 nm diameter) without targeting tags, we speculated that they would quickly disperse between cells within a tissue. We therefore tested in parallel QD800 nanoparticles functionalized with a peptide that facilitates its uptake into cells (referred to here as QD800.2; **Fig. 2a-b**).

**Figure 2.**
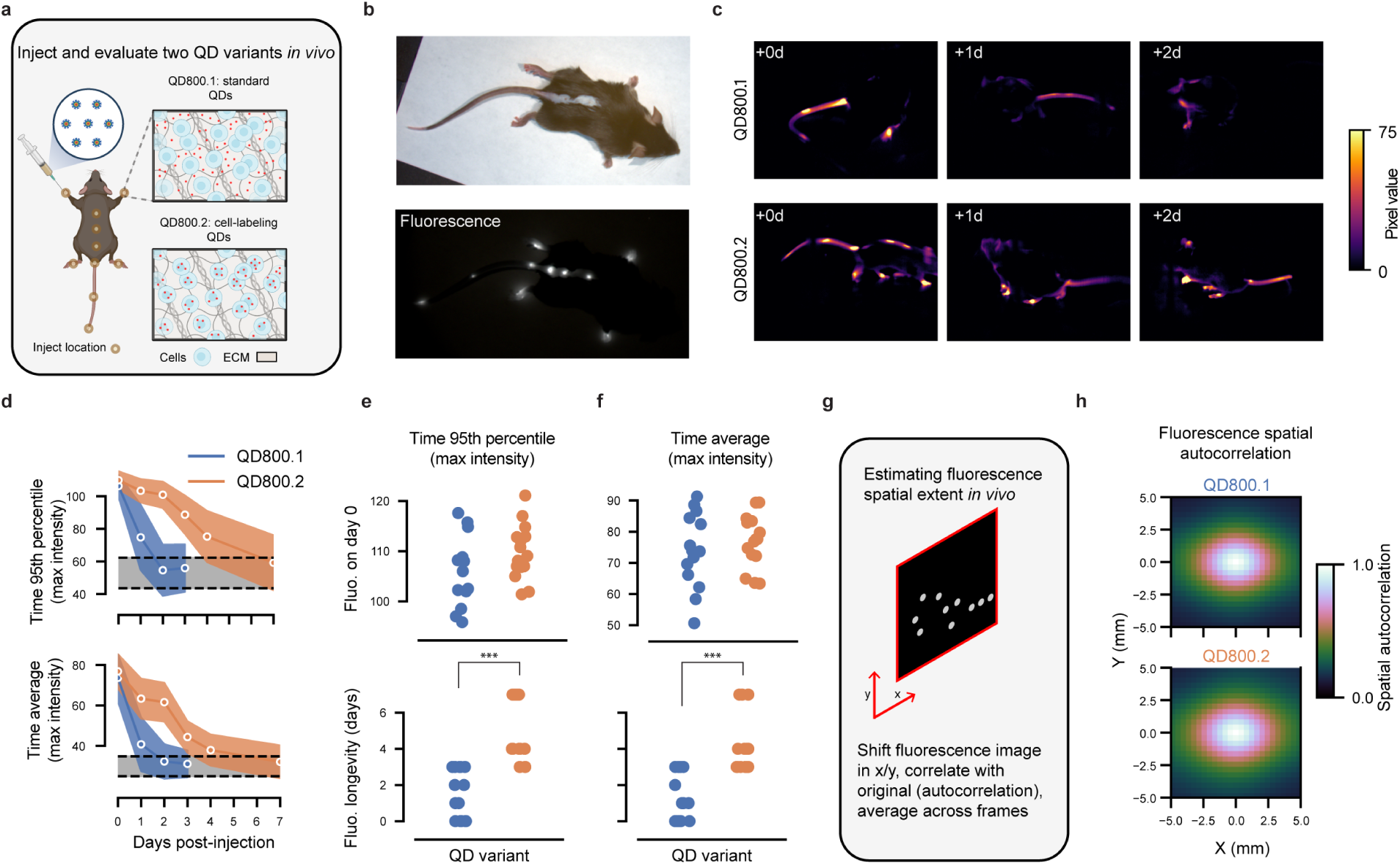
Half-life of QDs in buffer (QD800.1) and QDs that enter cells (QD800.2). a) Schematic representation of QD injection sites with QD800.1 and QD800.2 and their hypothesized aggregation in tissue. QD800.1 likely resides in the extracellular matrix, while QD800.2 is designed by the manufacturer to enter cells. b) Reflectance and fluorescence image of a mouse immediately after injection of QD800.2. c) Example fluorescence images from QD800.1 (top) and QD800.2 (bottom) imaged at 0, 1, and 2 days post-injection. d) Either 95^th^ percentile (top) or average pixel intensity (bottom) plotted as a function of days post injection for both variants (see Fig. 1f for schematic of quantification). Shown is the mean (line) and one standard deviation (shaded region) across each quantity for every mouse/camera view pair (n=15). Gray region indicates 95^th^ percentile confidence interval for vehicle/blank mice. e) Initial brightness (top) and the fluorescence longevity (time for the trace for each mouse/camera pair to cross below the 95^th^ percentile of vehicle/blank mice, bottom) for both variants calculated from 95^th^ percentile pixel intensity over time (p=.23, U=142, f=.63 for brightness; p=6.9e-6, U=219, f=.97 for longevity, Mann-Whitney U test; n=15 mouse/camera pairs each). f) Same as Fig. 2e, except computed using average pixel intensity over time (p=.41, U=133, f=.59 for brightness; p=7.3e-5, U=204, f=.91, for longevity, Mann-Whitney U test; n=15 mouse/camera pairs each). g) Schematic of spatial autocorrelation calculation to measure the length-scale of QD induced fluorescence. h) Average in vivo spatial correlation of fluorescence across all mice and camera views for both QD800.1 (top) and QD800.2 (bottom).

Longitudinal imaging of mice injected with QD800.1 and QD800.2 revealed that signal decayed within 1.6 ± 1.4 days and 5.3 ± 1.7 days, respectively (**Fig. 2d-f,** see **Extended Data Fig. 3-4 for raw data**). This supported our hypothesis that cell internalization could prolong visualization of the optical tag. Since QD800.1 and QD800.2 both use the same underlying fluorescent nanoparticle, it also suggested that the decay in fluorescence was not due to photobleaching. Average spatial autocorrelation of the fluorescence images across all animals and camera views showed that both QD800 variants remained localized to the injection site *in vivo* immediately following injection (**Fig. 2g-h**).

### Microbead-immobilized QDs yield a bright fluorescent spot that can be imaged for weeks

Even though QD800.2 outlasted QD800.1, both were only detectable for days, not weeks. We hypothesized that immobilizing QDs on large microparticles would further prolong the signal of the optical tag by minimizing diffusion and discouraging clearance from the body.

Prior work has shown that > 30 µm diameter microbeads injected subdermally in humans can form long-lasting deposits that persist for multiple years^47^. We, therefore, tested our hypothesis by combining biotin-functionalized QDs with streptavidin-functionalized microbeads (QD800.3; **Fig. 3a-e**) and injecting them subdermally as before. We tested six commercially available microbeads and found that porous approximately 85 µm diameter high-capacity agarose beads yielded bright beads (**Extended Data Fig. 5**). We confirmed that QD800.3 formed dense deposits in the skin that persisted in the animal (**Fig. 3f-g, Extended Data Fig. 6)**, increasing the imaging window to >16 days (16.6 ± 4.6 days; QD800.3, n=3 mice, **Fig. 4**). This longevity is critical to tracking the progression of skeletal kinematics in certain mouse disease models, such as models for Parkinson’s disease^48^, and over the course of standard systems neuroscience experiments that include reading and writing neural activity over multiple sessions.

**Figure 3.**
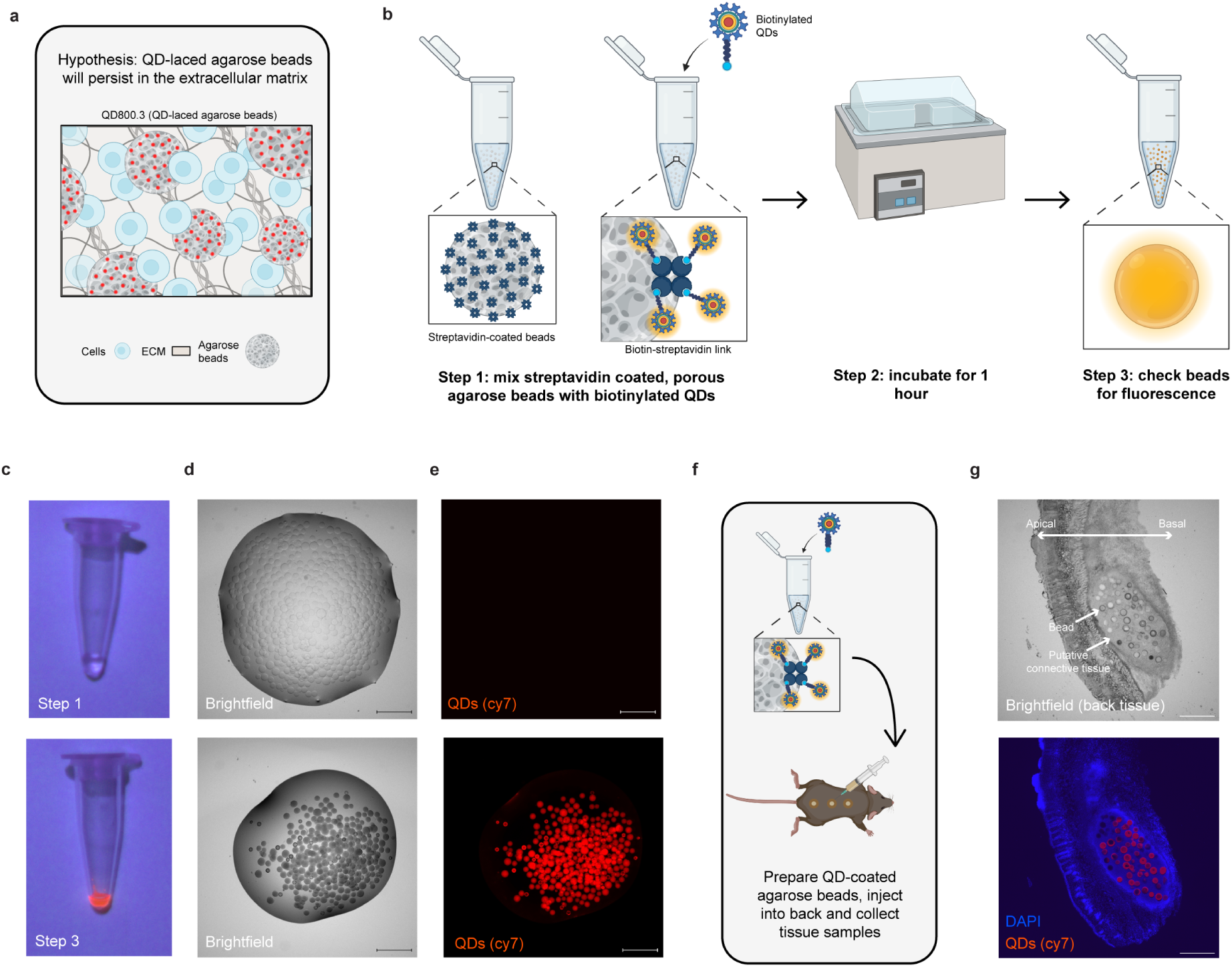
A method for attaching QDs to microscale, porous agarose beads using the streptavidin-biotin link. a) Schematic of our hypothesis – attaching QDs to a relatively large, biocompatible, porous agarose beads will lead to stabilization of QD fluorescence. We call this QD800.3. b) Protocol for QD800.3. Streptavidin-coated agarose beads are mixed with biotinylated QDs and incubated for 1 hour at 40°C, mixed halfway through the incubation period. The supernatant is removed, the solution is washed 3 times with 1X PBS, and the beads are resuspended in 2% sodium alginate. c) Images of agarose beads (top) and our agarose bead QD mixture (bottom). Images were taken under 405 nm illumination. d) A droplet of agarose beads (top) or the agarose bead QD mixture (bottom) under brightfield illumination. Scale bar represents 500 µm. e) Fluorescence from the same droplet using Cy7 excitation/emission (see **Methods** for details). Scale bar represents 500 µm. f) Schematic of the experiment, QD800.3 was injected into the back. Tissue samples were harvested and imaged. g) Brightfield (top) and fluorescence (bottom) images from the same tissue sample taken from the back 1 day after QD800.3 injection. Scale bar represents 500 µm.

**Figure 4.**
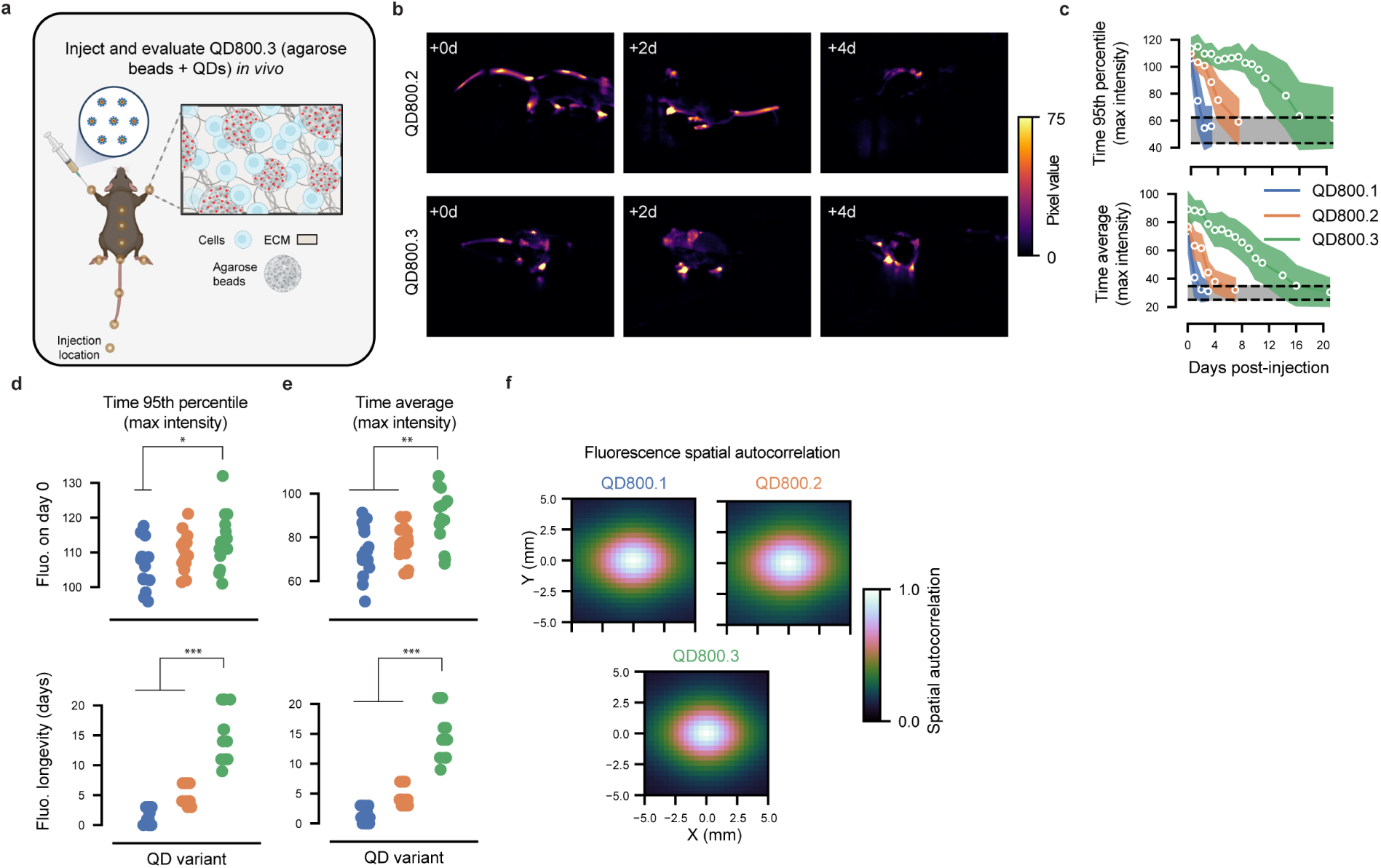
Custom agarose bead QD mixture (QD800.3) outperforms all other variants in vivo. a) Schematic representation of QD injection sites with QD800.3. b) Example fluorescence images from QD800.2 (top) and QD800.3 (bottom) imaged at 0, 2, and 4 days post-injection. c) Either 95^th^ percentile (top) or average (bottom) pixel intensities plotted as a function of days post-injection for all three variants. Line indicates the average and shaded region one standard deviation across mouse/camera pairs. d) Initial brightness (top) and the decay rate (bottom) for all variants computed using the 95^th^ percentile across time (p=.013/.19, U=173/144.5, f=.77/.64 for brightness; p=2.5e-6/2.3e-6, U=225/225, f=1/1 for longevity, Mann-Whitney U test; n=15 mouse/camera pairs each, QD800.3 compared with QD800.1/QD800.2). e) Same as Fig. 4d, except computed using the average across time (p=.004/.009, U=182/176, f=.81/.78 for brightness; p=2.6-e6/2.5e-6, U=225/225, f=1/1 for longevity, Mann-Whitney U test; n=15 mouse/camera pairs each, QD800.3 compared with QD800.1/QD800.2). f) Spatial autocorrelation for all three variants in vivo.

### Optimizing excitation wavelength and imaging optics maximizes signal-to-noise

Having confirmed the use of QDs as discrete, bright, long-lived optical tags, we next hypothesized that the signal-to-noise ratio (SNR) could be further improved by optimizing the imaging hardware (camera, excitation lights, and optical filters). Specifically, we speculated that tuning the excitation and emission optics to match peak QD800 excitation without being appreciably visible to the mouse (wavelengths equal to or above 650 nm), could substantially improve fluorescence signal^49^. Initial recordings were made using 730 nm LEDs with polarization filters for excitation, and cameras were equipped with 830 nm longpass filters and polarization filters tuned to the orthogonal orientation to minimize stray light. While 730 nm is ideal for penetrating the skin, it only excites QD800 particles at 1% efficiency, and an 830 nm longpass filter clips the peak emission (**Fig. 5a**)^33^. Hence, we altered our setup to use 660 nm LEDs, thereby doubling the excitation efficiency, and a 780 nm longpass to maximize collection of the emitted light (**Fig. 5a, Extended Data Fig. 7a**). These changes increased SNR > 2-fold for mice injected with QDs subdermally as before (**Fig. 5b-c**, **Extended Data Fig. 8**), which provided unambiguous detection of keypoints with noise characteristics suitable for use in downstream applications, such as training markerless keypoint models. We call this final iteration of QD800.3 tags combined with the upgraded imaging system and analysis pipeline, QD-Pi.

**Figure 5.**
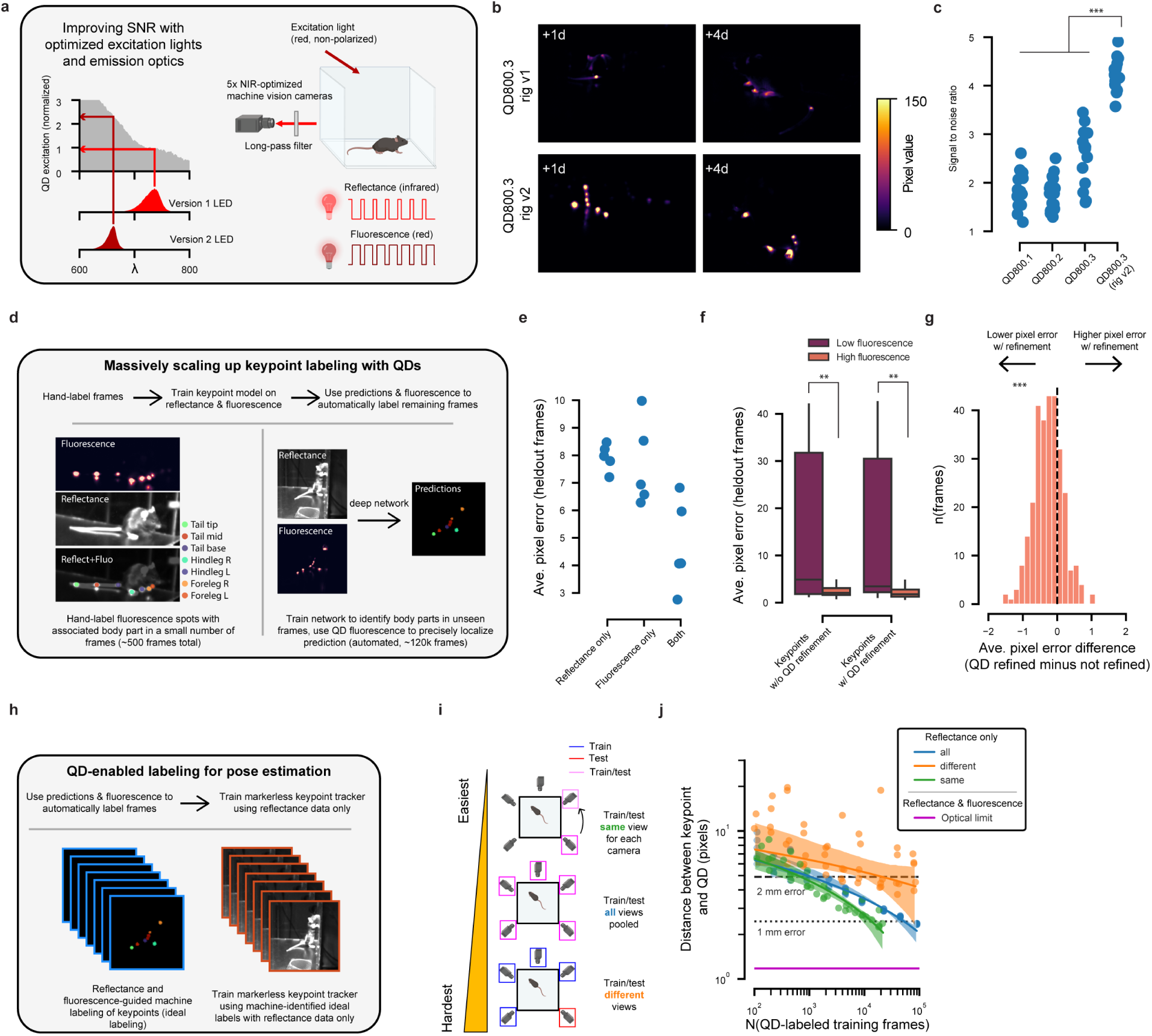
Combining QD800.3 and an optimized imaging rig enhances SNR, enabling both benchmarking and massive scaling of markerless keypoint trackers. a) Schematic of changes made to imaging rig to maximize SNR (also see **Methods**, **Extended Data Fig. 7**). b) Example fluorescence images from mice injected with QD800.3 using the first imaging rig (v1, top) or the second (v2, bottom). c) SNR across all variants and rigs, estimated by taking the standard deviation across x- and y-axes for each fluorescence frame and then averaging the result across all frames for each mouse/camera view pair (QD800.3 rig v2 comparison with all other variants, p=3.4e-6, U=0, f=1, Mann-Whitney U test, n=15 mouse/camera pairs). The noise level was estimated by calculating the same metric for blank/vehicle mice. d) Schematic illustrating massive scaling up of keypoint labeling using reflectance and fluorescence frames. Combined fluorescence and reflectance images were labeled by hand and used to train a U-Net to automatically identify body parts and corresponding fluorescence peaks in unlabeled data. The resulting predictions were then refined using the QD-based fluorescence data (see **Methods**). This resulted in 123,885 labeled frames, which we call QD-Pi-120K. e) Comparison of keypoint prediction performance (pixel error relative to hand-labeled ground-truth, L2 distance) using reflectance images alone (“reflectance only”), fluorescence images alone (“fluorescence only”), or the combination of the two (“both”). Each point is the average over all keypoints and frames for a model trained using a different random seed (n=5 restarts). f) Per-frame average pixel error for models trained using both fluorescence and reflectance data, computed separately for keypoints with high or low fluorescence (see **Methods**) (p=.004, U=1435, f=.29 without refinement; p=.0007, U=1257, f=.26 with refinement; n=290 high-fluorescence and n=17 low-fluorescence frames; Mann-Whitney U test). g) For each frame, the difference in pixel error with and without fluorescence refinement of the predicted keypoint (p=3.9e-21, W=7613, n=290 frames, r=- .64; Wilcoxon signed-rank test, see **Methods**). h) Schematic of our method for using machine-labeled frames to train markerless keypoint trackers on reflectance data only. i) Schematic of SLEAP training regimes. U-Nets were trained and tested using data from the same camera view (“same”, top), trained and tested using data from all camera views (“all”, middle), or trained on four camera views and tested on one heldout view (“different”, bottom). j) SLEAP was used to train U-Nets to identify keypoints in reflectance data using QD-Pi-120K according to the training regimes indicated in Fig. 5i. Error was considered the distance between each identified keypoint and the nearest QD fluorescence peak. The optical limit is the minimum feature size resolvable by our optical setup (see **Methods**). We consider 2 mm error minimum acceptable performance and submillimeter error necessary for tracking movement with similar resolution used in human movement studies. Note these error estimates are a lower bound for the expected error with human labeling, since the QD-labeled training sets are free from inter-annotator error. Lines indicate linear regressions fit on log-transformed data (random sample consensus regression), and shaded regions indicate 95% bootstrap confidence interval of the fit.

### Imaging datasets of QD800-tagged mice can be used to profile and train markerless keypoint tracking algorithms to reach new levels of precision and robustness

Having built QD-Pi, we set out to collect a unique, large-scale dataset that could be used to both train and benchmark markerless keypoint trackers. Specifically, we simultaneously collected reflectance and fluorescence images from freely moving mice injected with QD800.3 subdermally (n=3 additional mice), which enabled the collection of high-quality video-rate data surface features of the mouse registered to ground truth key points inside of the mouse (**Fig. 5d**).

This system allowed us to test the accuracy of commonly used markerless keypoint tracking algorithms relative to ground-truth optical markers placed inside of the body. Here, we focused on the commonly used U-Net architecture available in the SLEAP package^10^.

Our first goal was to automatically label the body part associated with each QD800.3 injection. To accomplish this, we hand-labeled 571 frames across the five camera views (see **Methods**). Fluorescence and reflectance frames were superimposed so that the labeler could identify both a given body part and peak QD800 fluorescence (**Fig. 5d**). These hand-labels were then used to train a U-Net using both reflectance and fluorescence data, to identify which QDs correspond to which body part. The resulting dataset comprised 123,885 frames of machine-labeled keypoints in three mice across five camera views, which we call QD-Pi-120K (see **Methods**). This presented us with a new opportunity to test the scaling properties of common markerless keypoint trackers relative to ground-truth fiducial markers (**Fig. 5d-g**). We speculated that such a large dataset would enable us to train individual networks that could identify keypoints across all five camera views, and networks that could generalize to novel views — a particularly challenging problem given the small size of hand-labeled datasets typical in keypoint tracking applications.

We experimented with training and testing U-Nets using frames from all camera views (“all”), training on four views and testing on the one held out view (“different”), and training and testing using frames from the same camera view (“same”) (**Fig. 5h-i**). Surprisingly, even when training and testing using frames from the same camera view, 700 frames were required to achieve a mean error of 2 mm. More frames were required for the other two scenarios: 1,100 using all camera views, and 21,300 when attempting to generalize to a novel view. To achieve submillimeter accuracy, we required 13,600 frames for the same camera view, and 52,600 frames using all camera views. When generalizing to a novel view, we were unable to achieve submillimeter accuracy with up to 100,000 frames (**Fig. 5j**).

### QD800 can directly track joint kinematics

Finally, we speculated that, since QD800.3 yielded bright, long-lasting discrete points of fluorescence under the skin, we could introduce QD800.3 directly into the knee joint and still acquire a resolvable signal. This would confirm that our new method can provide direct visualization of key body parts for precisely quantifying kinematics in mice.

Guided by an existing protocol^50^, we altered our QD injection protocol to allow for intra-articular injection of QD800.3 into both knee joints in a mouse. The first test was performed on a mouse cadaver (**Fig. 6a-b,** see **Methods**), followed by post-injection confirmation that movement of the overlaying skin did not affect the fluorescent spot. We further confirmed the location of the injection in relation to the skeleton with a micro computed tomography (microCT) image from a mouse cadaver using IVIS Spectrum CT (Perkin Elmer/Revvity; **Fig. 6c** and **Extended Data Video 2,** see **Methods**). Additionally, we performed intra-articular injections into the knee joints of live mice and subsequently imaged them in our plexiglass arena (1 hour post-injection, **Fig. 6d**). We confirmed that the intra-articular injection sites could be reliably tracked as the mice freely moved, with a signal level comparable to that of the subdermal injections.

**Figure 6.**
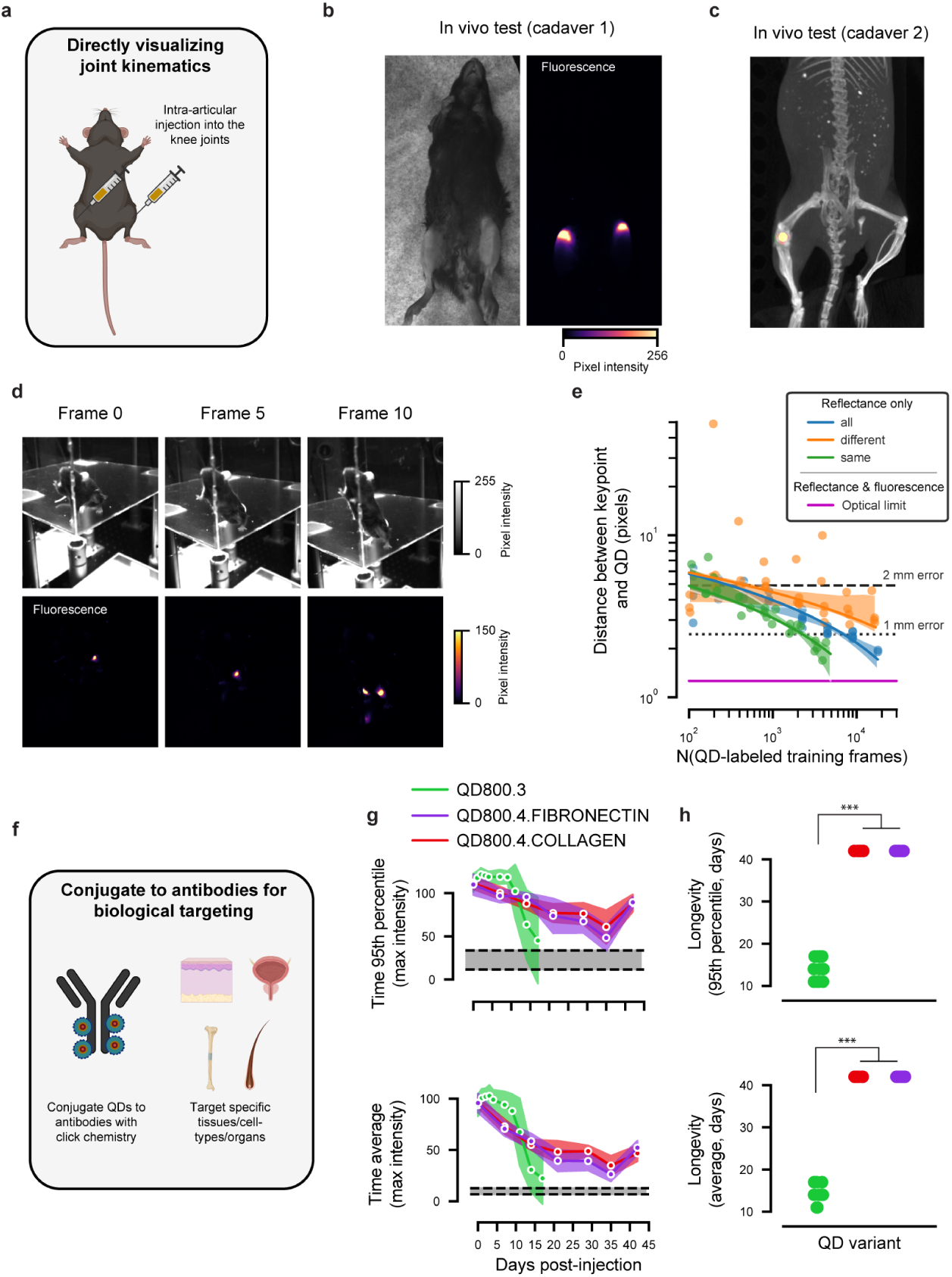
Direct visualization of the knee joint in vivo through intra-articular injection of QD800.3 and targeting to specific tissues with QD800.4. a) Schematic of experiment. QD800.3 was injected directly into the knee joints, then the skin and underlying fatty tissue above the injection site was moved with a pair of surgical tweezers to confirm the accurate targeting of the joint. b) In vivo cadaver validation of intra-articular targeting. c) Micro-computed tomography (microCT) image from the right knee joint of mouse cadaver. Fluorescence is overlaid (yellow spot) on top of a reconstructed mouse skeleton. d) Frames from video recorded 1 hour post-injection into live mice. A volume of 4 µL of QD800.3 was delivered to each knee joint using a micro-pipette. e) Using the same procedure as Fig 5j, we trained keypoint models to track knee joint location based on surface features. Plotting conventions are the same as Fig. 5j. f) Strategy for conjugating QD800 to antibodies in order to target specific tissues, cell types, and molecules. g) Fluorescence plotted against days post injection with same plotting convention as Figs. 2d and 4c. h) Quantification of longevity of fluorescence signal per mouse/camera pair using the 95^th^ percentile across time (p=1.5e-5, U=0, f=0 for longevity comparing QD800.3 to QD800.4.COLLAGEN, Mann-Whitney U test; p=1.5e-5, U=0, f=0 comparing QD800.3 to QD800.4.FIBRONECTIN; n=15 mouse/camera pairs for QD800.3, n=10 mouse/camera pairs for each QD800.4 variant, and n=10 vehicle-injected mouse/camera pairs) and using the average across time (p=1.3e-5, U=0, f=0 for longevity comparing QD800.3 to QD800.4.COLLAGEN, Mann-Whitney U test; p=1.3e-5, U=0, f=0 comparing QD800.3 to QD800.4.FIBRONECTIN).

Given the high SNR of QD800.3 directly injected into the knee joints, we hypothesized that a markerless keypoint tracker could be trained to accurately track the position of the joints. To test this, we again hand-labeled a small number of frames (n=822 from n=1 mouse) to map each fluorescent point to the corresponding body part (left or right knee joint). These labels were then used to train SLEAP to leverage reflectance and fluorescence data to label the full dataset. Finally, we used this dataset to benchmark the ability of SLEAP to track the position of the knee joints given surface features (**Fig. 6e**). Here, we found that 1 mm accuracy could be achieved using 2,300 frames for the same camera view, 7,000 frames for all camera views, and more than 15,000 frames (the maximum number tested) when generalizing to a novel view.

### QD800 can be conjugated to antibodies for targeting specific molecules, cell types, and tissues

Having established QD800.3’s enhanced performance in labeling adipose tissue and joints, we speculated that QDs could also be deployed in applications that required additional biological specificity; that is, QDs could be used in applications that required highlighting specific proteins, cell types, or tissues. As a proof of principle, we adapted a protocol for conjugating QDs to antibodies via click chemistry (see **Methods**). Here, we tested directly conjugating QDs to antibodies that target ultra-long-lived proteins in the extracellular matrix in skin and joints: collagen and fibronectin (QD800.4.COLLAGEN and QD800.4.FIBRONECTIN, **Fig. 6f-g**). Remarkably, we found that both QD800.4 variants outperformed QD800.3, with resolvable fluorescence up to 42 days post-injection.

## Discussion

Here we developed a new method called QD-Pi for optically tracking movement in freely moving mice. To achieve this, we injected NIR-I-emitting QDs (QD800), free or immobilized on microbeads, in discrete spots in the adipose tissue just beneath the dermis of each animal. We found that the microbead-immobilized particles formed long-lived deposits that can be used as fiducials for tracking keypoints inside of the body using standard machine vision cameras. We found that two commercially available variants were suitable for *in vivo* use: QDs originally designed for tracking the vasculature (QD800.1) and QDs tailored for cell labeling (QD800.2). However, the fluorescence of both variants decayed to noise levels within 1-5 days after injection. This motivated us to design a custom third variant where biotinylated QDs are attached to porous, streptavidin-functionalized agarose microbeads (QD800.3). QD800.3 led to robust labeling with fluorescence that remained resolvable for over 2 weeks after injection, a timescale compatible with standard systems neuroscience experiments. Lastly, we show that QDs can be used to directly visualize joints, a key step in accurately resolving kinematics in mice, and customized for biological targeting.

QDs represent an attractive alternative to other tags employed in marker-based methods that have been used in rodents. Elegant work from Butler et al. demonstrated that UV ink can be applied to the fur and used to measure a single point in freely moving mice^43^ (see also^51^ for a similar method using non-fluorescent paint applied to rodents). However, UV excitation light may be visible to the animal^49^, the ink can fade, and similar markings have been shown to be relatively imprecise^16^. Since the ink is applied to the outer layer of the skin and fur, it can also be groomed or licked off by the mouse. Other studies have used piercings with highly reflective markers that can be tracked using commercial motion capture systems originally designed for use in humans^16,44^. However, these markers are bulky and still represent motions on the surface of the skin. Another marker-based work from Moukarzel et al. applied tattoo ink directly to the tissue surrounding the knee joint of rats^45^. Although this method targets the musculoskeletal system, it appears to have relatively low SNR since the tattoo ink is simply dark under IR illumination rather than fluorescent (we note that SNR was not quantified), the ink likely disperses widely around the joint preventing precise localization, and it has only been shown to mark a single joint at a time.

As an alternative to marker-based methods or X-rays, markerless keypoint trackers have capitalized on recent developments in image-based deep learning methods, resulting in a surge of interest in movement tracking^5–10^. However, they often require laborious hand-labeling of training data, which leads to imprecise predictions due to inter-labeled jitter and ambiguity in labeling certain keypoints^13,16^. Our method enabled the creation of a massive dataset with “ground-truth” labels for keypoint trackers — that is, labels with well-defined physical positions inside of the body — which we call QD-Pi-120K. Using the resulting dataset, we found that only hundreds of frames were required to achieve an acceptable threshold of 2 mm average error with markerless keypoint trackers, in line with prior work^10,43^. However, human kinematics studies typically track kinematics with submillimeter precision, and humans are on the order of 20 times larger than laboratory mice, and thus the relative error of kinematic tracking in mice is still much higher relative to studies of movement in humans. Achieving submillimeter average error or less in our hands requires at least 13,000 labeled frames, with at least 50,000 required if one wants a single network that works across multiple views, or more than 100,000 to generalize to a novel view. Datasets of this size are not feasible with manual labeling, which emphasizes the critical need for techniques like ours that enable the collection of large high-quality datasets for training the next generation of markerless keypoint trackers. Moreover, in addition to training markerless keypoint trackers, QDs can be used directly for high-resolution kinematic tracking without these steep requirements on the amount of training data.

Despite the impressive performance and ease-of-use of markerless keypoint trackers, these methods fundamentally rely on surface features of the animal. Prior X-ray videography of freely moving rodents has shown that, even if one can optimally track keypoints on the skin, they can substantially distort the movement of the underlying joint^21^. Our method is a promising approach to accurately measure the part of the body under more direct control of motor circuits — the muscles, joints and bone — and more comprehensively map the relationship between neural activity and movement. We show that this direction is feasible, as QDs can be injected into the knee joint for direct tracking of joint kinematics.

However, while using QDs as optical tags is an attractive approach, they still have certain limitations. Signals from QD800.1 and QD800.2 rapidly faded with a timescale of 1 and 5 days, respectively. Prior work has speculated that QD800.2 may quickly leak from cells either through excretion or cell division, which could explain our observations^52,53^. Additionally, resolving signal from QD800 injections in the fatty tissue along the back required shaving the fur. Finally, despite the radically improved longevity of QD800.3 and QD800.4 variants, we aim to achieve lifelong labeling in the future, which may be possible given the photostability of QDs.

Another limitation is the mechanism of QD clearance from the body. It has been reported that QDs that were administered intravenously and intradermally get deposited in the liver and lymphatic and renal systems of mice and rats^46,54,55^. The mechanism of clearance of microbead-immobilized QDs, as used here, remains an open question and needs to be further investigated. An additional limitation is the speed of our current multiplexed imaging system, 30 frames per second, with approximately 60 milliseconds between the reflectance and fluorescence exposures. Thus, there is the possibility of minor displacements between the fluorescence data and surface features in QD-Pi-120K during periods of rapid motion. Subsequent versions of the imaging rig will focus on the time between these exposures to minimize the possibility of this occurring.

Future directions include injecting QDs directly into the muscle, bone, and specific tissues within a joint such as the tendons. Another direction is to label other tissues like the whiskers, and potentially internal organs including the bladder. Highly specific targeting to cell-types, tissues, and molecules will be enabled by further variations on QD800.4. Moreover, QDs have near-ideal optical properties for spectral multiplexing. QDs are available at a wide variety of emission wavelengths spanning most of the visible spectrum into mid-IR with high quantum yields^56,57^, with conveniently overlapping excitation spectra. Consequently, one could easily label separate mice or separate parts of the body with distinct colors.

Thus, we present a new method for directly tracking positions inside of an animal’s body. In addition to advancing the measurement of kinematics in freely moving mice, we envision that our method can be used to augment markerless keypoint trackers, which have become the standard method for movement tracking in laboratory animals^5–10^. By imaging both the fluorescence from specific points in the body and the surface features surrounding those fluorescent points simultaneously, we have amassed a large dataset that can be used to benchmark models for tracking keypoints in 2D^5,7,9,10^ and 3D^6,8^. Moreover, in addition to acting as a method for directly measuring the motion of positions inside of a mouse’s body, our method will be instrumental in building ground-truth datasets at the scale necessary for constructing foundation models for tracking motion in laboratory animals. Ultimately, this method can be used to resolve complex motor patterns in rodent models of movement disorders with newfound accuracy and precision.

## Acknowledgements

We are grateful to members of the Markowitz lab for assistance in labeling videos for training markerless keypoint trackers. Lisette Bahena assisted with preliminary experiments. Dr. Alan Emanuel provided guidance on performing and interpreting histology of the skin. Dr. Laxminarayanan Krishnan provided guidance on IVIS Spectrum CT imaging system. Dr. Timothy Gardner provided useful discussion at the onset of the project. Drs. Anqi Wu, Ariel Levine, Caleb Weinreb, Lena Ting, and Young-hui Chang provided useful comments on the manuscript. The work was supported by a Career Award at the Scientific Interface from the Burroughs Wellcome Fund (JEM), the David and Lucille Packard Foundation (JEM), and the Sloan Foundation (JEM). Certain illustrations presented in this study were created using BioRender.com (Fig. 1a-d, Fig. 2a, Fig. 3a-b, f, Fig. 4a, Fig. 5a, i, Fig. 6a,f, Ext. Data Fig. 6).

## Author Contributions

J.E.M. and E.Z.U. originally conceived of the use of quantum dots as fiducial markers. J.E.M., E.Z.U., A.P., and D.K. conceived of the use of microbeads for increasing longevity. J.E.M., E.Z.U., A.P., and D.K. designed experiments. E.Z.U. and A.P. performed experiments. E.Z.U. performed histology. J.E.M. and E.Z.U. analyzed the data and created figures. J.E.M., E.Z.U., A.P., and D.K. wrote the manuscript.

## Methods

We used quantum dot (QD) solutions that were suitable for *in vivo* use. For this manuscript, we used four variants of QDs, all of which have a CdSe core and ZnS shell and are additionally coated with polyethylene glycol (PEG) for enhanced biocompatibility (QD800.2 is functionalized with polyarginine peptide instead of PEG).

### QD800 Variant 1 (QD800.1)

Due to their prior use in *in vivo* tracing of the vasculature in mice, the first variant we tried was QTracker^TM^ 800 Vascular Labels (ThermoFisher Scientific, #Q21071MP). These were used undiluted (2 µM).

### QD800 Variant 2 (QD800.2)

Due to their prior use in *in vivo* tracking inside of cells, the second variant we tried was QTracker^TM^ 800 Cell Labeling Kit (ThermoFisher Scientific, #Q25071MP). Qtracker® nanocrystals (Component A, 2 µM) was diluted by half with Qtracker® carrier (Component B).

### QD800 Variant 3 (QD800.3)

Both QD800.1 and QD800.2 were relatively short-lived in the skin. Larger particles, e.g. polymethyl-methacrylate (PMMA) beads, can persist in the skin for up to 5 years^47^. Hence, we hypothesized that attaching the QDs to larger biocompatible particles would enhance longevity. We found using agarose beads to be ideal due to their porous structure, biodegradability, and biocompatibility. Moreover, we utilized the streptavidin-biotin reaction to immobilize the QDs onto the agarose beads.

High-capacity streptavidin agarose beads (Amid Biosciences, #SA-101-1) and custom-made biotin-conjugated QDs (ThermoFisher, per manufacturer’s specification, ≥0.5mg/mL) were used for agarose bead injections (see **Extended Data Fig. 5** for all microbead brand names that were tested). For suspension, sodium alginate (Sigma-Aldrich, #W201502-SAMPLE) was hydrolyzed in 1X phosphate-buffered saline (PBS) to make a 2% (w/v) solution. The beads were spun down, and the supernatant was removed. QD solution was added at 2:1 (v:v) beads-to-quantum-dots ratio and gently mixed with the beads by pipetting up and down. The mixture was left to incubate at 40°C for 1 hour and was mixed halfway through the incubation period to ensure saturation. After the incubation, the solution was mixed again and then washed three times with 1X PBS. The supernatant was removed, and the beads were resuspended in 2% sodium alginate in a 1:1 beads-to-alginate ratio to allow for an even distribution during injection.

### QD800 Variant 4 (QD800.4)

To enable biological targeting of QDs, we used antibody-based targeting of QD800 to long-lived ECM-associated proteins, such as collagen I and fibronectin. The exact concentration of QDs used in this kit was not disclosed by the manufacturer.

To conjugate QD800 to anti-Collagen I (Abcam, #ab34710) and anti-fibronectin (Abcam, #ab2413) rabbit polyclonal antibodies, we capitalized on an enzyme- and click chemistry-mediated site-specific antibody conjugation approach^2^ and used an antibody labeling kit (Invitrogen, SiteClick^TM^ Antibody Labeling Kit, #S10455). Anti-collagen I-QD800 and anti-fibronectin-QD800 antibody conjugates are referred to as QD800.4.COLLAGEN and QD800.4.FIBRONECTIN, respectively. Briefly, the antibody was concentrated using membrane filtration. Next, the carbohydrate domain of the antibody was modified for azide attachment using uridine-5′-diphosphate-N-Azidoacetylgalactosamine (UDP-GalNAz) labeling. Azide-modified antibody was then concentrated using membrane filtration and conjugated to DIBO-labeled QD800. To enhance the yield and concentration of conjugated antibodies (QD800.4) and to filter out unconjugated antibodies, we purified QD800-conjugated antibodies using membrane filtration.

### Surgical Procedure

All procedures were approved by the Georgia Institute of Technology Institutional Animal Care and Use Committee (IACUC; protocol #A100557). Twenty-two male and nine female adult (13-20 weeks old) C57BL/6J mice were purchased from the Jackson Laboratories and kept under a reverse 12h light/12h dark cycle with *ad libitum* access to food and water. Specifically, n=3 mice were not injected with any material (blank), n=5 were injected with buffer (vehicle, n=3 were used for rig v1 and n=2 for rig v2. Blank and vehicle were combined in the control group), n=3 were injected with QD800.1, n=5 with QD800.2 (n=3 were used for rig v1 and n=2 for rig v2), n=6 with QD800.3 (n=3 were used for rig v1 and n=3 for rig v2), n=3 were injected with QD800.3 intra-articularly into the knee joints, n= 2 were injected with QD800.4.FIBRONECTIN, and another n=2 were injected with QD800.4.COLLAGEN. For **Fig. 2b**, n=1 mouse was injected with QD800.2 (sex was not determined for this mouse). The animals were anesthetized with 4% isoflurane mixed in air using a nose cone. The procedures were carried out under 1.8-2% isoflurane anesthesia on a heating pad and completed within 30 minutes. The fur along the spine was shaved and further treated with a hair removal cream. The QD mixtures were injected using a sterile pulled glass micro-pipette (Drummond 3-000-210-G) created using a Sutter P-2000 laser puller (parameters: heat 450, filament 4, velocity 150, delay 175, pull 35). The micro-pipette was attached to a modified positive-displacement microinjector (Drummond 3-000-510-X). Specifically, the stainless-steel plunger of the microinjector was cut to enable the pulled micropipette to be safely attached to the microinjector. The QD mixtures were delivered to each of the 14 injection sites subdermally: the paws (dorsal and ventral), the tail (base, midsection, and tip), and the back (upper, middle, and lower midline dorsal region). Following each injection, 70% ethanol and triple antibiotic ointment were applied to minimize potential infections. The animals were allowed to recover in their cage for one hour after the procedure.

A volume of 2 µL of QD800.1 and QD800.2 was delivered to the 14 injection sites subdermally with approximately 0.1 mm diameter micro-pipette. For agarose bead injections, the diameter of the micro-pipette was approximately 0.3 mm to allow for the passage of the beads. The injection volume was changed to 4 µL to maximize the signal and was delivered to the same 14 injection sites subdermally.

An injection volume of 4 µL was also used to deliver QD800.4.FIBRONECTIN and QD800.4.COLLAGEN to the same 14 injection sites subdermally with approximately 0.1 mm diameter micro-pipette. The vehicle and QD800.2 were repeated with the same injection volume alongside these experiments. Additionally, the fur along the spine was shaved and treated with a hair removal cream under anesthesia 4 and 6 weeks after QD800.4 injections to enhance the keypoint signal from the back of the mice.

The protocol for intra-articular knee injection was based on the protocol from Pitcher et al. paper and modified accordingly^50^. A volume of 4 µL of QD800.3 was delivered to each knee joint using micro-pipette with a diameter of approximately 0.2 mm to both allow for bead passage and accurate access to the joint. For *in vivo* injections, post-operative analgesic was given at the start of the procedure, and following each injection, 70% ethanol and triple antibiotic ointment were applied. The animals were allowed to recover for one hour after the procedure. Here, n=3 mice were injected, and 2 out of 3 mice showed strong fluorescence in only one knee joint. The third mouse showed strong fluorescence in both knee joints and was thus used for analysis in **Fig. 6d-e**.

### Recording Arena

A plexiglass arena was created from clear cast-acrylic panels (McMaster-Carr, #8560K184) to form an open-top cube with 29.845 cm edge length (i.e. same width, depth and height). Interdigitated patterns were cut into the edges of panels to ease fitting together. Panels were glued together using acrylic plastic cement (Sci-Grip, #10315). Custom 3D-printed molds were secured to the bottom acrylic panel (Torr-Seal Epoxy, Varian), and screw-to-expand brass inserts were inserted into the molds to secure ¼-20” set screws. The other end of the set screws was inserted into Thorlabs 1” optical posts to, in turn, secure to an optical breadboard placed on top of a leveled frame (PFM52503).

### Recording Sessions

Mice were placed into the plexiglass chamber and imaged for approximately 3-5 minutes per session.

### Recording Hardware – Version 1

The plexiglass arena was filmed using five hardware-synchronized NIR-optimized Basler USB3 cameras (acA2040-90um). For wide field-of-view imaging, the cameras were outfit with a Thorlabs machine vision lens with an 8 mm focal length (MVL8M1). In order to prevent imaging of QD excitation light, long-pass (MidOpt LP830-55) and polarization filters (PR1000-55) were secured to the lens. Polarization filters were rotated until excitation light was minimized. For exciting QDs at wavelengths compatible with imaging through skin we used NIR-I emitting LED lights (SL246-730IC) outfit with polarization filters (PA371-S82). For collecting reflectance images, IR-emitting LED lights were used (SL246-850IC). The lights were triggered in a temporally multiplexed configuration so that fluorescence and reflectance data could collected near-simultaneously using the following sequence: (1) IR lights on for 10 milliseconds (2) all lights off for 1 millisecond (3) NIR lights on for 23 milliseconds, (4) all lights off for 1.5 milliseconds. Power to the LED lights was supplied from a benchtop voltage source (BCK Precision BK1550). Lights were triggered using 5V signals generated from an Arduino Uno, which was also used to trigger camera exposures via the GPIO lines on the Basler cameras. The illumination sequence was repeated for approximately 5 minutes for each recording session. In order to capture mice at all angles, the cameras were arranged in a pentagonal formation surrounding the plexiglass arena.

### Recording Hardware – Version 2

The following modifications were made to the recording hardware version 1 (see above section) to maximize SNR. First, in order to enhance the excitation of the QD800 nanoparticles, we decided to slightly blue-shift the excitation light (**Fig. 5a**). The 730 nm LED lights were replaced with 660 nm LED lights (Advanced Illumination SL-S100150W-660). Due to the blue-shifted excitation light, polarizing filters were no longer needed to filter out stray light, so they were removed to enhance signal levels. Additionally, the long-pass filters were modified to accommodate the new wavelength, so they were replaced with 780 nm long-pass filters (MidOpt LP780-55).

### Recording Hardware – Optical Limit

The optical limit of our system, presented in **Figs. 5-6**, was estimated by calculating the Nyquist limit given a pixel size of 5.7 µm, an object distance of 304.8 mm (the approximate distance from each camera to the center of the plexiglass arena), and a lens focal length of 8 mm.

### Recording Software

Camera control and image acquisition software was written in Python. Custom software was also written for the Arduino to trigger the cameras over the appropriate GPIO line along with the LED lights.

### Analysis of Quantum Dot Fluorescence

To assess the brightness and longevity of QD injections, we computed a simple summary metric for each camera view and each session (**Fig. 1f**). First, to remove any contribution of the background, a rolling background (1500 frame sliding window, non-overlapping) was subtracted from the fluorescence frames (**Extended Data Fig. 2**). Next, to summarize the intensity on a per-frame basis, we computed the maximum pixel value across x and y for every frame. Finally, to summarize either the average or peak intensity across time for each camera view and each recording session, we computed either the mean or the 95^th^ percentile across maximum frame intensities.

### Markerless Keypoint Tracking

To track keypoints, we utilized SLEAP^10^. First, reflectance and fluorescence frames from all five camera views were alpha blended (90% fluorescence and 10% reflectance), uploaded to segments.ai, and were hand-labeled. Hand-labels were verified by a second labeler. Next, the hand-labeled, blended frames were used to train a U-Net to automatically identify body parts that coincided with fluorescence peaks. 571 frames from all five cameras were hand-labeled using the segments.ai platform. We found the following U-Net parameters in SLEAP yielded reasonable performance in validation data: filters 64, filters_rate 1.75, middle_block true, up_interpolate true, max_stride 64, stem_stride NULL. The following augmentation settings were used (any options not listed were set to false): rotate_min_angle −15, rotate_max_angle +15, translate_min −50, transate_max +50, scale_min .85, scale_max 1.15, gaussian_noise_mean 5.0, gaussian_noise_stddev 1.0, contrast_min_gamma .5, contrast_max_gamma 2.0, random_flip true, random_flip_horizontal true. This network was then applied to our entire dataset. The predicted x/y position of each body part was then corrected using the fluorescence channel. In order to precisely localize each keypoint prediction, for each prediction we computed the fluorescence center of mass within a 7.5 pixel radius surrounding the predicted keypoint. The fluorescence center of mass distance from ground-truth labels was compared with raw keypoint predictions in **Fig. 5f-g.** We refer to fluorescence localization as refinement. The center of mass was then used as the “ground-truth” x/y position for training and assessing the markerless keypoint trackers shown in **Fig. 5h-j**.

As a rule of thumb, given an approximately 10 mm span between many key landmarks on a mouse^5^, we assume that the average error of a given keypoint tracker should not exceed 2 mm, which is approximately 4.9 pixels given our optical setup (**see Methods**). A more stringent criterion of submillimeter error is likely required for accurately quantifying rodent kinematics (2.45 pixels on our setup), since submillimeter precision is common in human motion capture, and humans are approximately 20 times larger than mice^6–8^. Here, we define error as the average L2 distance between the network-identified keypoint and the corresponding QD fluorescence peak.

### Histology

The tissues of interest at the injection sites (back skin, paws, and tail) were harvested and kept in 4% PFA solution for 48-72h at 4C before transferring them to 1X PBS solution. Prior to cryopreserving with sucrose, the paws and the tail were further dissected to collect the skin at the injection site. The tissues were placed in 15% and then 30% sucrose in 1X PBS until they sunk. The tissues were embedded in OCT (Sakura Finetek, #4583) and frozen with dry ice. They were then cryosectioned at 50 µm, collected with microscopy slides, and stored at −80°C until further processing. The sections were thawed, washed twice with 1X PBS, incubated in DAPI (1 µg/ml in water) (ThermoFisher, #D1306) for 3-5 minutes, and washed thrice with 1X PBS. The slides were then coverslipped using an anti-fade mounting medium (VectaShield, #H-1700).

### Imaging

Sections were imaged with a Nikon Eclipse Ti2 equipped with a Lumencor Spectra III light engine and a Hamamatsu Orca Flash 3.0 CMOS sensor. Images for brightfield, DAPI (365 nm excitation, 10% laser power, 10 ms exposure time, 12-bit sensitive mode), and CY7 (730 nm excitation, 25% laser power, 10 ms exposure time, 16-bit mode) were collected at 4X and 10X magnifications with a Semrock filter cube (LED-DA/FI/TR/Cy5/Cy7-A-000).

### IVIS Spectrum CT imaging

Intra-articular knee injection of QD800.3 was performed to both knees of a mouse cadaver. Bottom half of the mouse was shaved and further treated with hair removal cream to reduce imaging artifacts.

IVIS Spectrum CT Imaging system (Perkin Elmer/Revvity) was used to image the right knee in Fluorescence Tomography (FLIT) mode using a set of trans-illumination points combined with CT based reconstruction of the skeletal structure. Living Image 4.7.4 software was used to reconstruct the fluorescence light source (QD800) based on this trans-illumination at selected locations. The reconstructed fluorescence was then overlayed with both surface reconstruction of the animal and its skeletal structure to create a 3D representation of the injected QD800.3. Only results from the right knee are shown since the left knee imaging parameters were not appropriately optimized for this demonstration.

### Statistics

All hypothesis tests were two-tailed and non-parametric. Where appropriate, we list the exact *p*-value, sample size (along with definition of a sample), and the exact value of the relevant test statistic. Results are expressed as mean ± standard deviation. In all figures, * denotes *p*<.05, \*\**p*<.01, and \*\*\**p*<.001. Effect sizes for Mann-Whitney U tests are given as common language effect sizes, f. Effect sizes for Wilcoxon signed-rank tests are given as the rank-biserial correlation, r.

---

**Extended Data Figure 1.**
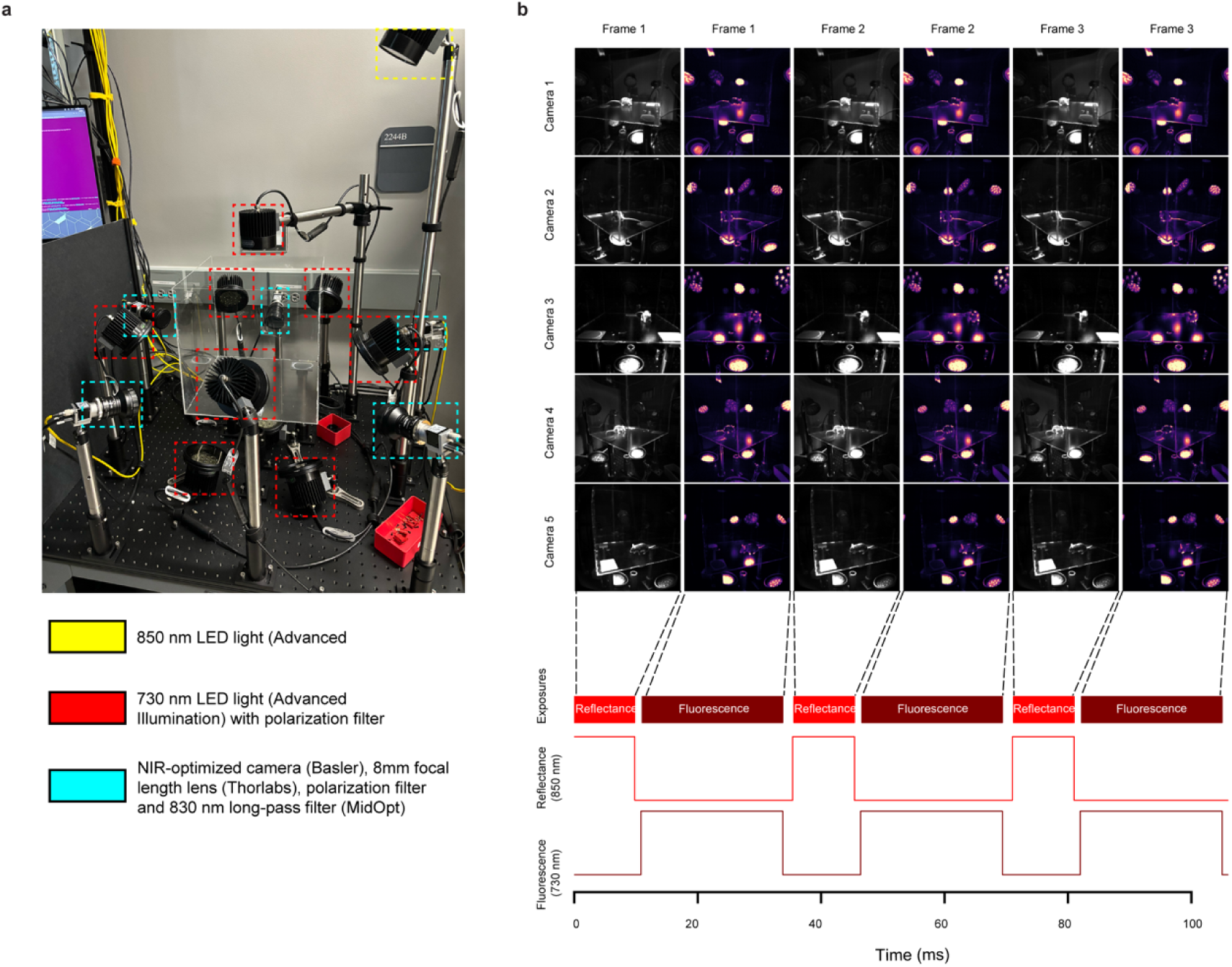
QD imaging rig version 1. a) Shown is the rig and plexiglass arena used for imaging QDs in freely moving mice. All data included in **Figs 1-4** and **Extended Data. Figs 2-4** were collected using this setup. b) Schematic of the illumination sequence used for temporal-division multiplexing.

**Extended Data Figure 2.**
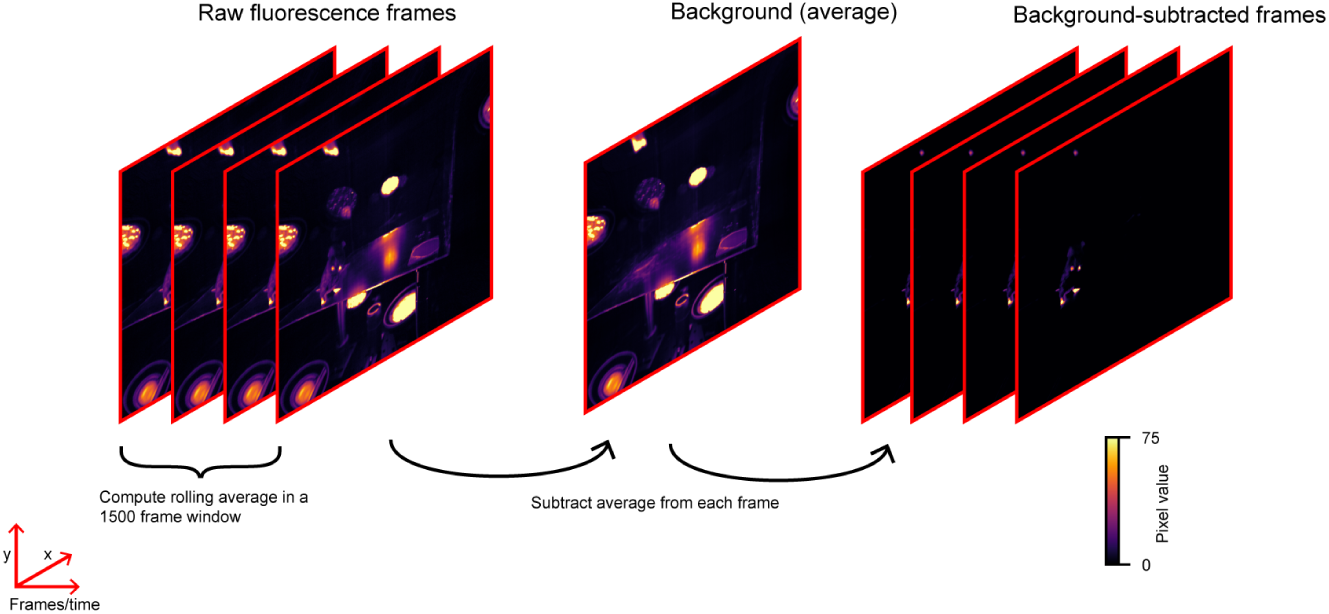
Fluorescence data pre-processing. To remove static elements from the scene (*e.g.*, excitation LEDs), the background of the image was computed using a non-overlapping 1500-frame-long sliding window. All visualization and quantification of fluorescence data uses background-subtracted data.

**Extended Data Figure 3.**
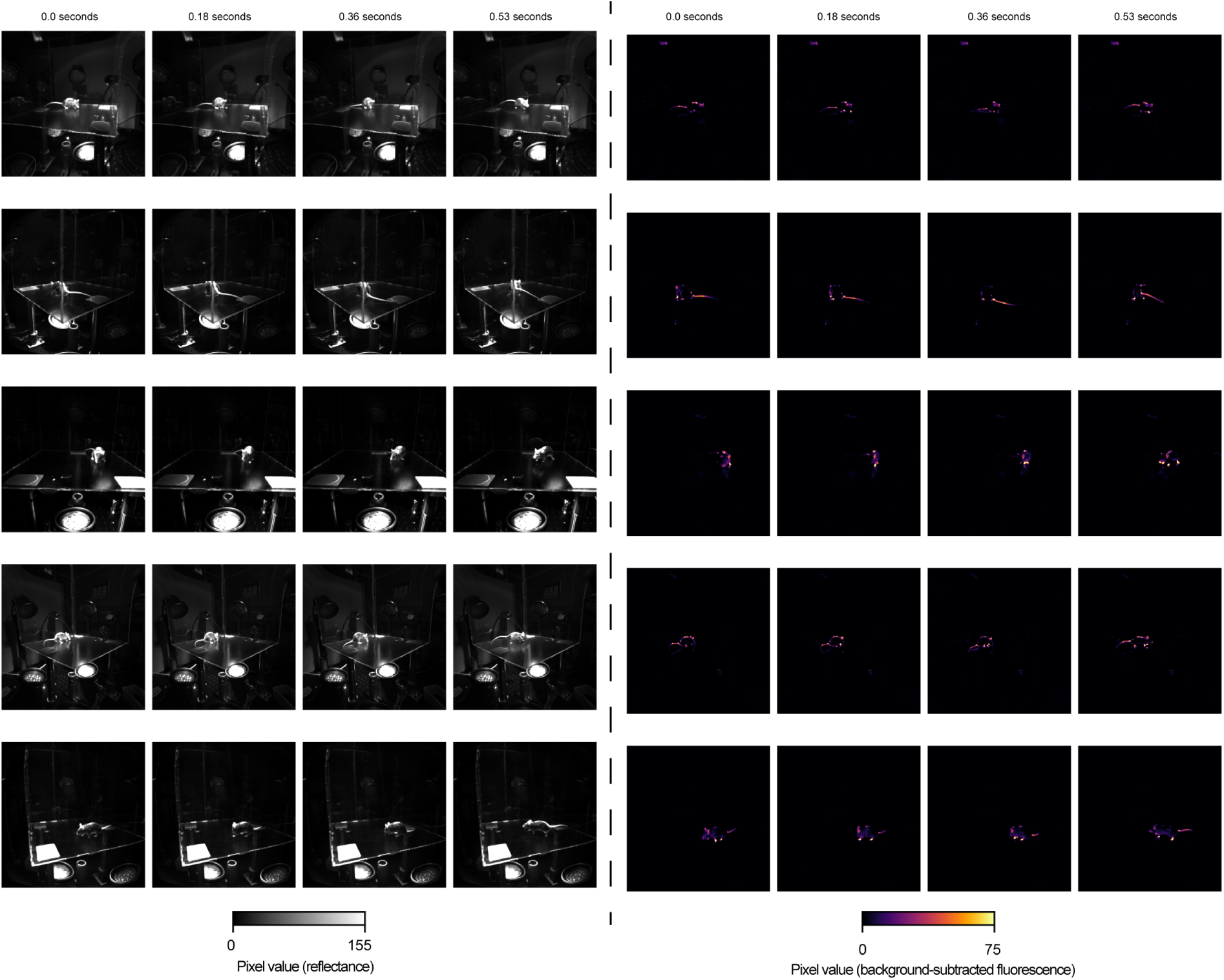
Example reffectance and ffuorescence data from a single session. Shown are reflectance (left) and fluorescence data (right) for four frames – timestamps are given at the top – from all five hardware-synchronized cameras. Data from a mouse injected with QD800.2.

**Extended Data Figure 4.**
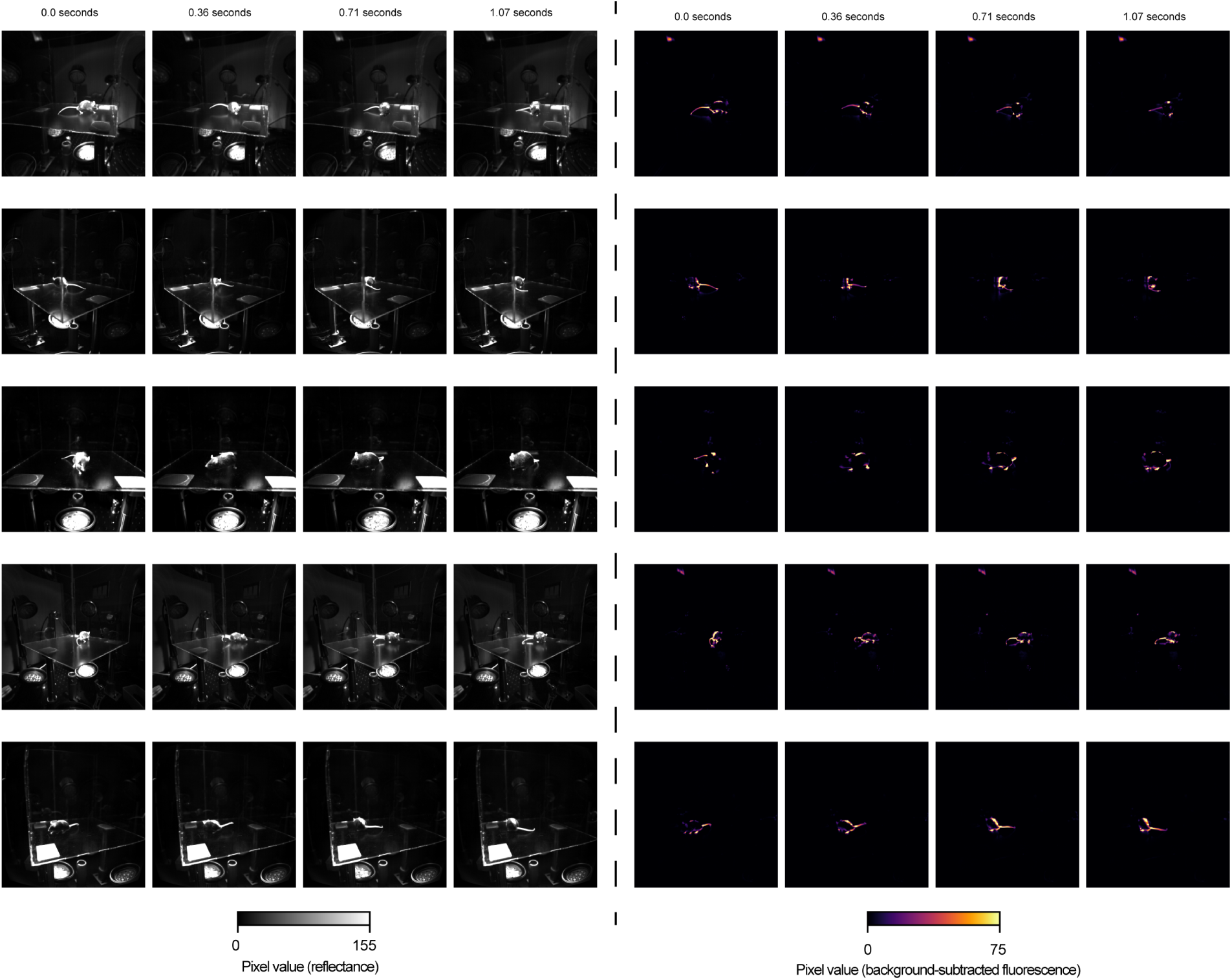
Example reffectance and ffuorescence data from a single session. Same layout as **Extended Data Fig. 3**, except a different session is shown. Data from a mouse injected with QD800.2. The mouse shown here is different from the mouse shown in **Extended Data Fig. 3**.

**Extended Data Figure 5.**
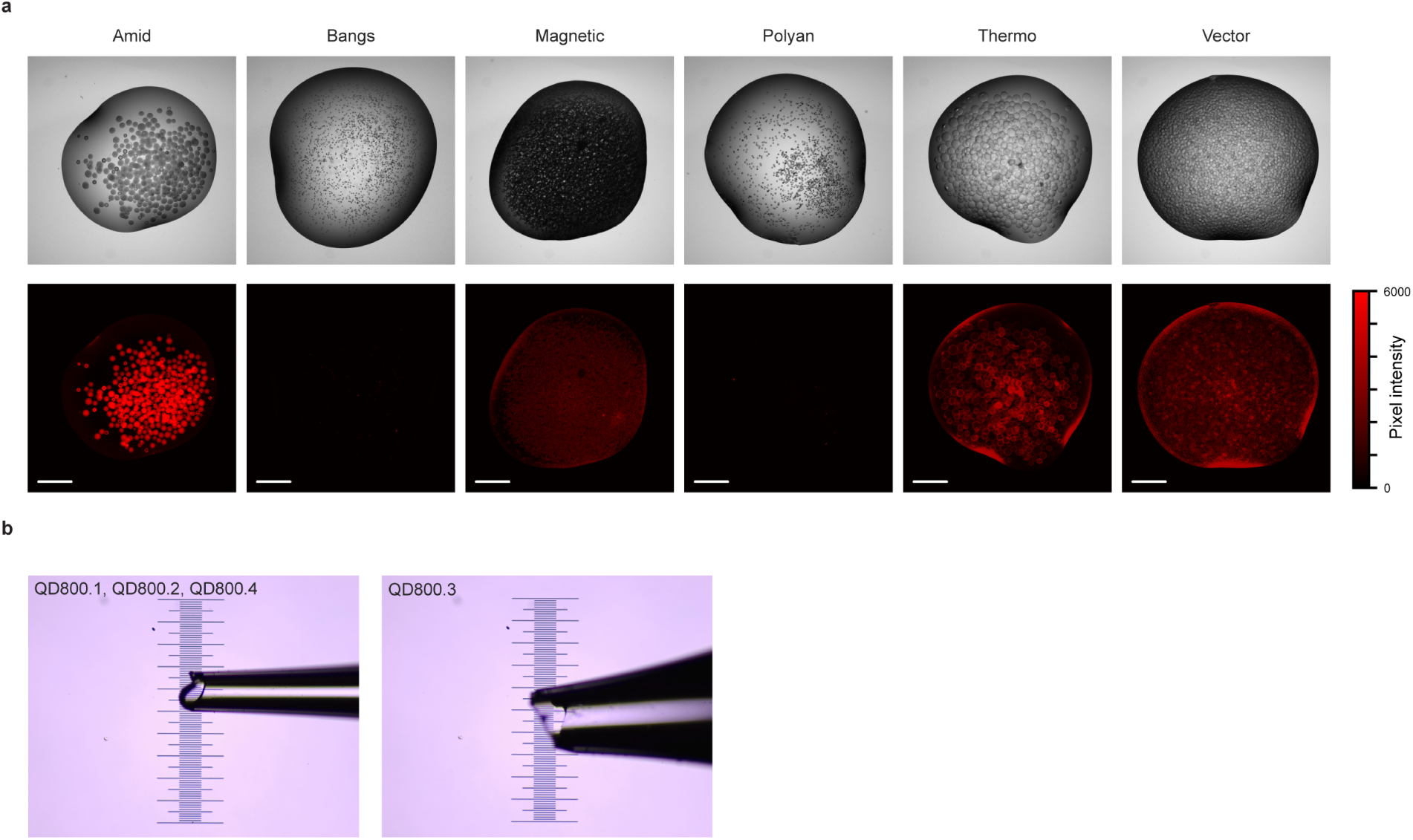
Testing different brands of microbeads. a) A droplet from six different microbead brands (Amid Biosciences, #SA-101-1; Bangs Laboratories, #CP01008; Resyn Biosciences, #MR-STM002; PolyAn, #105-21-020; Thermo Scientific, #20357; Vector Laboratories, #N-1000-002) mixed with biotinylated QDs imaged under brightfield (top) and fluorescence (bottom) using Cy7 excitation/emission (see **Methods** for details). Scale bar represents 500 µm. b) Pipette tips for QD injections. Each minor division is 10 µm.

**Extended Data Figure 6.**
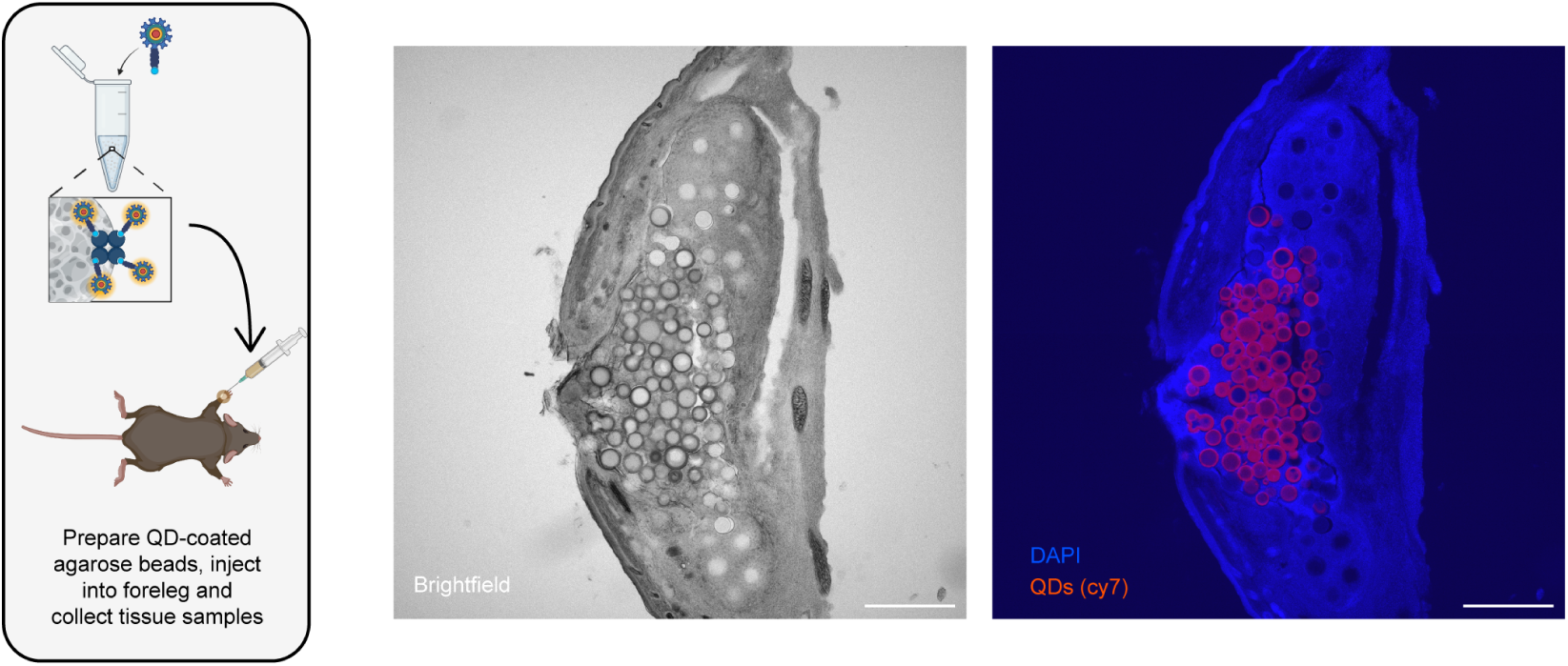
Histology from a left forepaw injection of QD800.3 (agarose bead QD mixture). Left, schematic of experiment. QD800.3 was injected into the left forepaw, then tissue was harvested 1 week post-injection and imaged. Middle, brightfield image of injection site. Right, fluorescence image of injection site. Scale bar represents 500 µm.

**Extended Data Figure 7.**
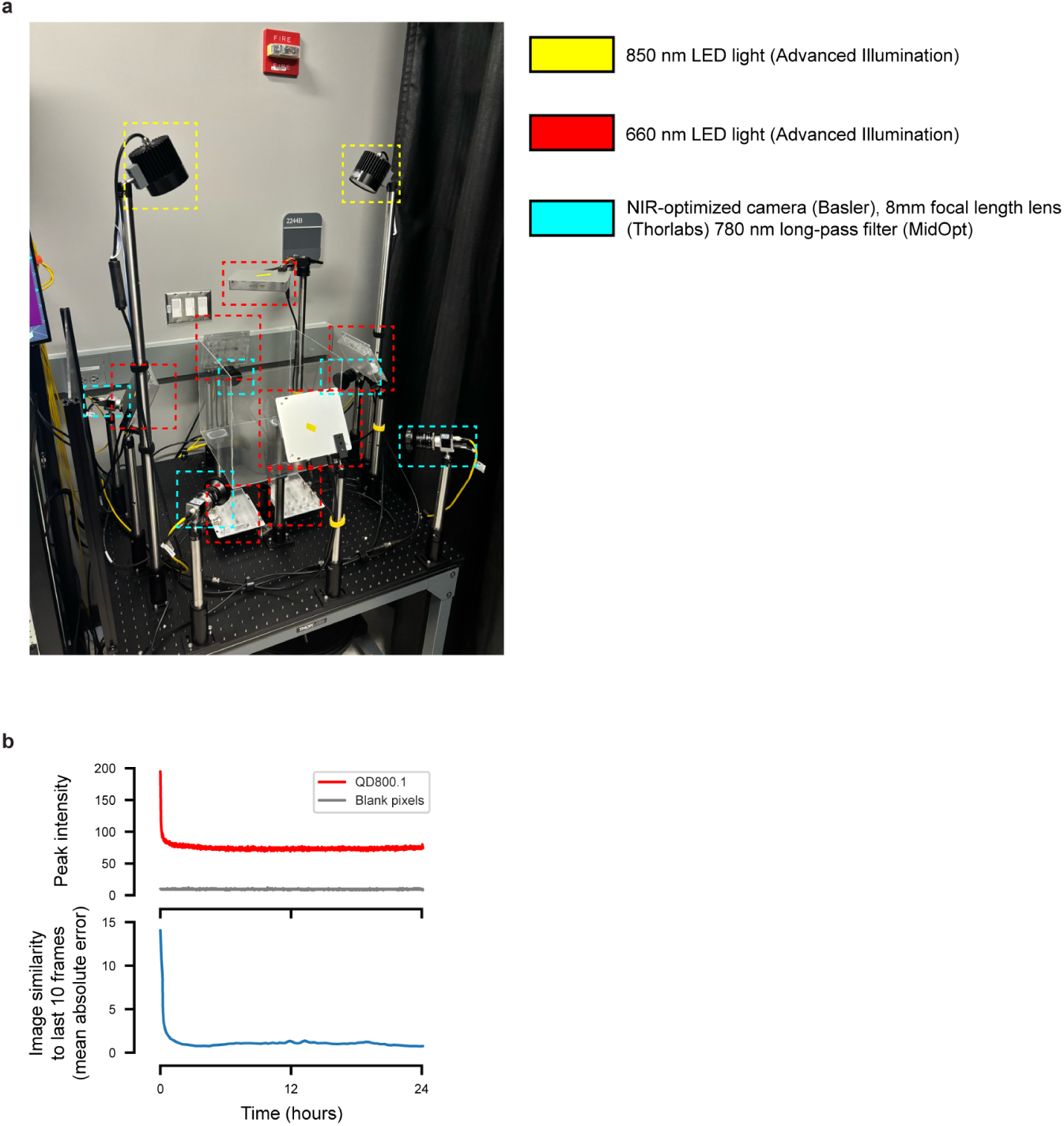
QD imaging rig version 2. a) All data included in **Figs 5-6, Extended Data Video 1**, and **Extended Data Fig. 8** was collected using this setup. b) QDs were pipetted onto a slide and imaged in the plexiglass arena once every 30 seconds for 24 hours in order to measure photostability in rig version 2. Excitation light was left on continuously. Top, an ROI was selected around the slide, and the peak was computed over the ROI per frame. Shown is the peak intensity in the image ROI and the peak intensity of a blank patch of the same frames. Bottom, similarity of the ROI to an average of the last ten frames. Due to drying, the QD droplet rapidly changes shape over the first hour. Once the spatial distribution of fluorescence is stable, the QDs are highly photostable.

**Extended Data Figure 8.**
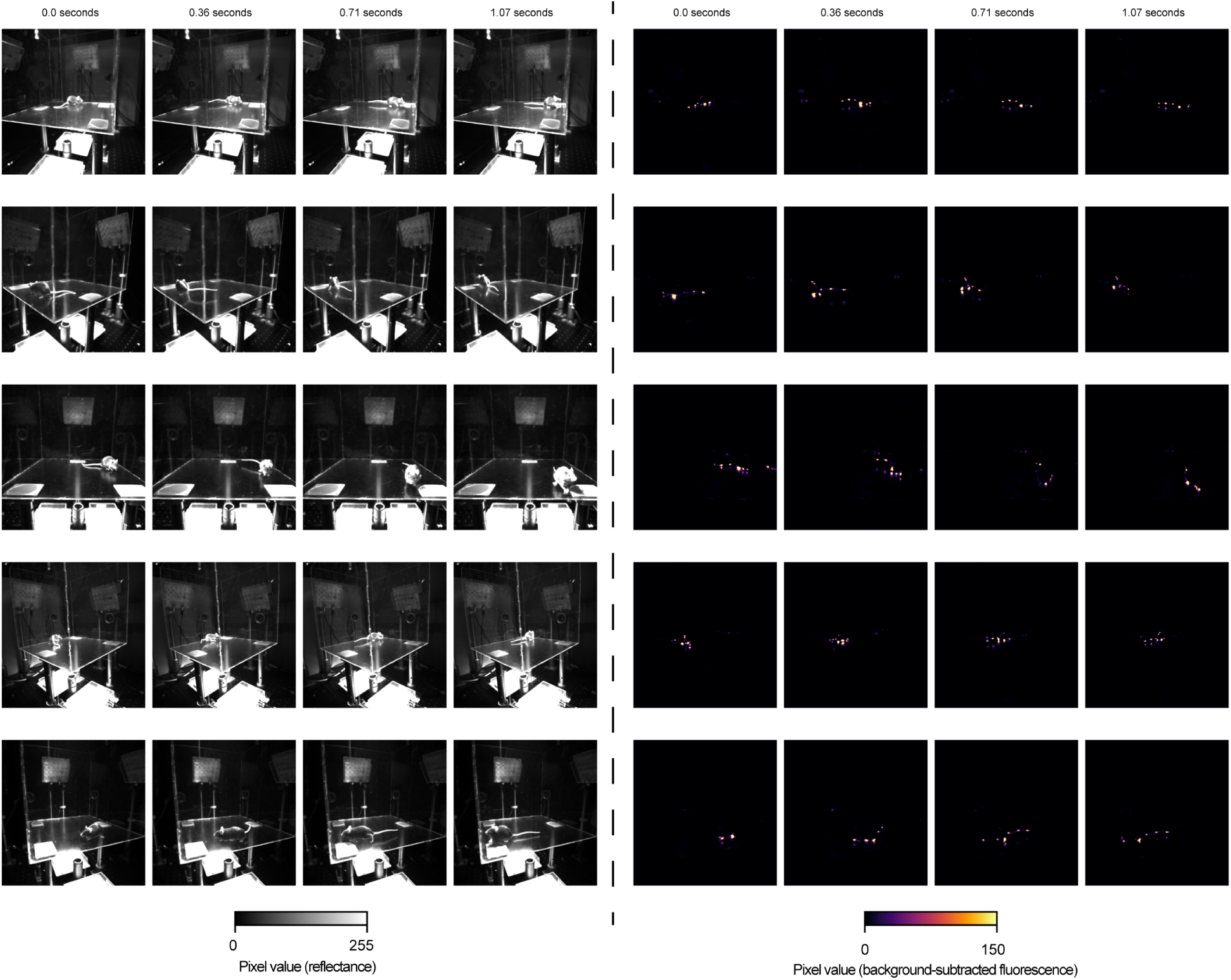
Example reffectance and ffuorescence data from a single session collected using rig version 2. Same layout as **Extended Data Figs. 3, 4**. The mouse shown here was injected with QD800.3.

***Extended Data Video 1. Example reffectance and ffuorescence video from all camera views from a single session.*** Shown is data from a single session with QD800.3, imaging rig v2.

***Extended Data Video 2. Series of X-ray images of mouse legs overlayed with ffuorescence in transaxial plane.*** A scroll through the transaxial plane to demonstrate localization of the fluorescent signal with respect to the right knee joint.

